# Efficient Inference, Training, and Fine-tuning of Protein Language Models

**DOI:** 10.1101/2024.10.22.619563

**Authors:** Muhammed Hasan Çelik, Xiaohui Xie

## Abstract

Protein language models have shown great promise in predicting protein structure, function, and the effects of missense variants on protein fitness. However, their use has been limited by the substantial computational resources required. In this work, we focus on improving the computational efficiency of protein language models (PLMs), specifically the Evolutionary Scale Modeling (ESM) family, to increase the accessibility of PLMs. By implementing optimizations such as FlashAttention and Partition-Attention, a novel technique designed to handle proteins of variable length, we achieved a 16-fold speedup in inference time and reduced memory usage by 3 to 14 times for long proteins. Additionally, 4-bit quantization applied to billion-parameter models led to a 2 to 3 times reduction in memory consumption with minimal performance loss in the missense variant effect prediction task. Training efficiency was also improved, with a 6-fold reduction in runtime achieved through activation checkpointing and the DeepSpeed Zero-Offload strategy. For fine-tuning, we employed parameter-efficient methods, enabling state-of-the-art predictions of protein properties and functions by training only the model head or a small fraction of adapter weights. For instance, we achieved a Spearman’s correlation coefficient of 70% in melting point prediction and an 87% area under the precision-recall curve (AU-PRC) for transcription factor prediction. Our efficient ESM (ESME) implementation significantly lowers the barrier to using these powerful models, making them accessible to academic laboratories with limited computational resources. The ESME implementation is available on PyPI (pypi.org/project/esm-efficient) and GitHub (github.com/uci-cbcl/esm-efficient).

## Introduction

An in-depth understanding of protein properties and function is crucial for the discovery of the mechanisms underlying essential cellular processes [1]. Deep learning techniques have long been employed for various in-silico protein prediction tasks [2] and recently led to breakthroughs in structure prediction [3], [4], [5], [6]. Furthermore, PLMs enable the prediction of protein properties [7], [8], function [9], and the effect of the missense variants on protein fitness and disease [10], [11], [12]. Recent advances have been achieved by the adaptation of large language models (LLMs) based on the transformer architecture from natural language processing (NLP) [13], [14] to the proteomics field [8], [15], [16], [17], [18], [19]. These models, known as protein language models (PLMs), leverage abundant protein sequence data and are trained using semi-supervised objectives to predict masked amino acids from the surrounding protein sequence context [20] [21] (Figure 1A). As a result, the attention mechanism in the transformer layers captures the complex interactions between the amino acids during the semi-supervised training to predict masked amino acids. In contrast to traditional sequence-based alignment, which only captures pairwise conservation between amino acids, deep learning models capture high-order epistatic interaction and context-dependent sequence constraints [22], [23]. Thus, PLMs can perform a wide range of downstream tasks, including protein structure prediction [3], [6].

**Figure 1:**
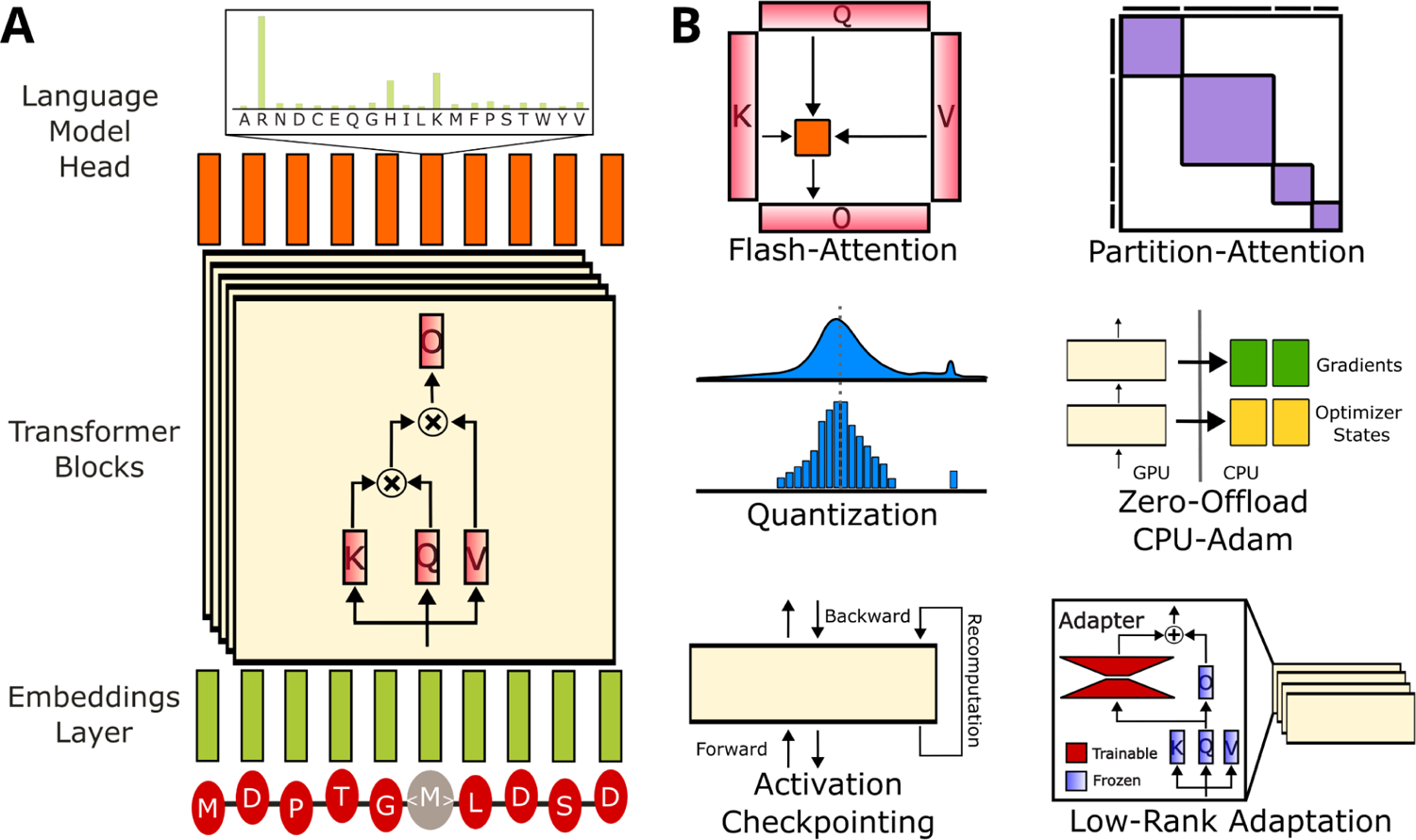
Overview of the protein language model architecture and efficient large language model techniques. **(A)** The architecture of the protein language models consists of three components: embedding layer, transformers block, and language modeling head. The embedding layer transforms amino acids into vector representations while encoding their positions within the sequence. The transformer blocks employ a self-attention mechanism to capture interactions between residues. Finally, the language model head predicts the likelihood of each amino acid. **(B)** The efficient LLM methods utilized to optimize inference, training, and fine-tuning of protein language models include FlashAttention, Partition-Attention, quantization, activation checkpointing, Zero-Offload/CPU-Adam, and Low-Rank Adaptation (LoRA).

Protein language models (PLMs) facilitate protein prediction tasks, particularly when supervised training data is limited. These models are fine-tuned to predict biochemical properties by leveraging predicted structures and contextualized sequence embeddings. For example, after fine-tuning with a small number of additional training samples, PLMs can accurately predict various biochemical properties, such as folding stability [21], [24], [25] and melting point [26].

By capturing intrinsic features and conserved amino acids, PLMs enable effective generalization through few-shot learning. Beyond structural predictions, PLMs also infer biophysical features, such as protein dynamics in disordered regions [27] and phase transitions [28]. Since the properties of proteins determine their functions, PLMs are also highly effective in predicting the functional characteristics of proteins. For instance, PLMs can accurately predict Gene Ontology (GO) terms [9] and Enzyme Commission (EC) numbers [29], [30] directly from amino acid sequences. Furthermore, the prediction of protein-protein and protein-small molecule interactions [5] is crucial for understanding the role of proteins in biological processes, and PLMs can predict these interactions, including ligand-binding proteins [31], peptide-T cell receptor interactions [32], and drug-protein interactions [33], [34], [35]. Finally, PLMs have important applications in protein design, enabling the generation of amino acid sequences with specified characteristics [36], [37], thereby facilitating novel approaches in in-silico drug development.

Another application of protein language models is the variant effect prediction of missense variants [10], [11], [12]. Classifying deleterious missense variants from benign ones is critical for the identification of pathogenic variants in disease studies. The PLMs learn the natural sequence space of the proteins selected by evolution; thus, they can estimate the variants disrupting the protein fitness. Learning the sequence distribution of homolog proteins and the co-conservation of amino acids reveals the epistatic interactions and improves variant effect prediction [22], [23], [38].

A significant challenge in utilizing protein language models (PLMs) is their substantial computational cost, with training requiring thousands of GPU hours on large computing clusters [39]. This high computational cost makes training PLMs from scratch impractical, especially for researchers relying on academic laboratory-grade computation infrastructure, thereby limiting the ability to experiment with different modeling choices and hindering progress in research.

Furthermore, inference and fine-tuning of these models are computationally expensive; our initial experiments revealed that inference on long protein sequences often results in memory errors. Fine-tuning, which requires backpropagation, further increases memory requirements. The large memory footprint also constrains the batch size during fine-tuning, which is essential for stable training and objectives like contrastive learning. Recent advances in efficient inference [40], fine-tuning [41], and training methods [42], [43] in the LLMs literature offer a promising solution to these limitations; thus, adapting these techniques could also facilitate the broader use of protein language models.

In this work, we implemented techniques to enhance the efficiency of protein language models and benchmarked their performance across various tasks, including variant effect prediction, using our reimplementation of the widely used ESM family, including ESM1v, ESM1b, and ESM2 models. We integrated widely used techniques like Flash-Attention [44] and introduced a novel approach called Partition-Attention to efficiently handle proteins of variable lengths, leading to a substantial reduction in both inference and training time, as well as memory usage. These reimplemented models achieve performance comparable to the original models in missense variant effect prediction, confirming the correctness of our implementation. Moreover, we achieve state-of-the-art performance on protein properties and function prediction by only training a small fraction of model weights. Overall, the approaches employed in this work greatly improve the efficiency of protein language models and enable their use on academic computing infrastructure. Our open-source efficient implementation ESME is available on PyPI (pypi.org/project/esm-efficient), GitHub (github.com/uci-cbcl/esm-efficient), and HuggingFace (huggingface.co/mhcelik/esm-efficient).

## Results

We applied a combination of efficient LLM techniques to the Evolutionary Scale Modeling (ESM) models, chosen for their standard transformer architecture, which facilitates the easy adaptation of LLM methods from NLP. We conducted in-depth benchmarking of the original ESM and efficient ESME reimplementation for inference, training, and downstream fine-tuning tasks (Figure 1B).

### Benchmarking the Model Inference

The first technique we leveraged in the efficient ESME implementation is FlashAttention [44]. FlashAttention is an efficient, IO-aware self-attention algorithm that optimizes memory usage and runtime for long sequences by minimizing read/write operations between HBM and SRAM while fusing operations to reduce memory overhead (Methods). To demonstrate performance gains, we sampled proteins of varying lengths from UniProtKB/Swiss-Prot [45] and measured runtime and memory usage. Inference performance was benchmarked on NVIDIA A100 GPUs using ESM2 models ranging from 8 million to 15 billion parameters, with a batch size of 16 proteins in brain floating point (bfloat16) [46] for both original and efficient implementations.

ESM1v and ESM1b were not included in this benchmark, as their behavior is comparable to the similarly sized ESM2 650M model. Adapting FlashAttention can reduce model runtime by ∼3 to 10 times across all model sizes, depending on protein length (Figure 2A, Table S1). For example, inference on a batch of 16 proteins with lengths of 300–400 residues takes ∼0.07 seconds with the ESME-2 650M model, compared to ∼0.21 seconds with the original implementation. The runtime of the models with FlashAttention scales linearly with input size, whereas the dot-product attention in the original implementation scales quadratically. Similarly, memory usage scales quadratically with input size in the dot-product attention implementation, limiting the batch size and inference on large proteins (Figure 2B, Table S2). Long protein sequences are often truncated to avoid an out-of-memory (OOM) error; however, FlashAttention maintains a small memory footprint even for large proteins. For instance, the 3-billion-parameter ESME-2 model uses approximately 5.6 gigabytes (GB) of GPU memory for a batch of proteins 300–400 amino acids long and no more than ∼8.4 GB for proteins up to 3,500 amino acids. In contrast, the original implementation cannot handle proteins longer than 3,000 amino acids for billion parameter models.

**Figure 2:**
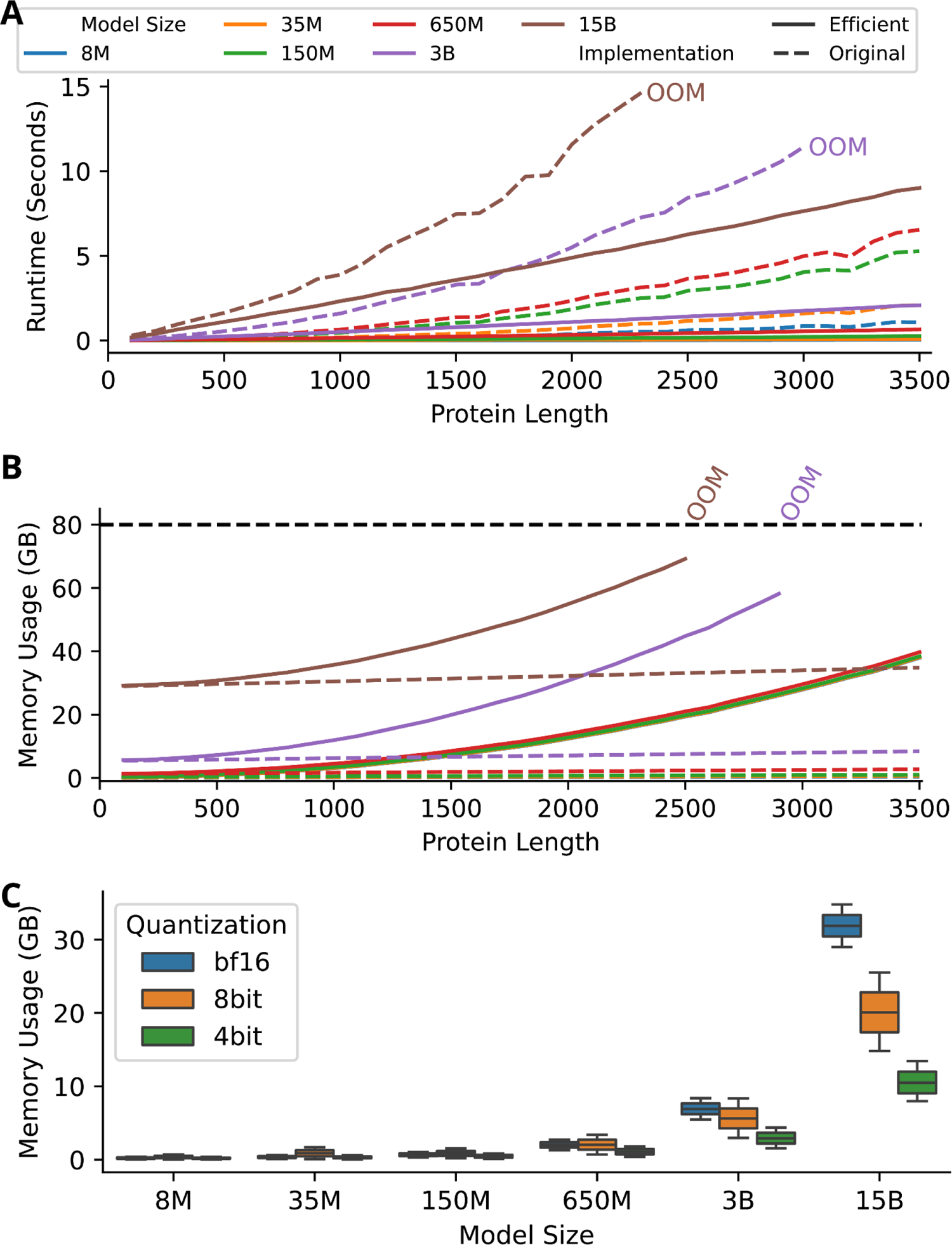
The benchmark of inference performance. **(A)** The runtime comparison between the original ESM2 model (solid line) and ESME2 (dashed line) demonstrates the reduction in runtime as protein length increases. M and B are abbreviations for million and billion, respectively. **(B)** The benchmark of memory usage shows that FlashAttention scales linearly with sequence length, while dot-product attention scales quadratically. **(C)** The model quantization techniques effectively reduce memory usage, but their impact is significant only in billion-scale models.

The proteins vary significantly in length; for example, ∼12% of human proteins are longer than 1000 amino acids, while the median protein is 414 amino acids long (Figure S1.A, B). With linear scaling, GPU memory usage of a batch of proteins can be quantified by token (amino-acid) count in the batch. Consequently, we set batch sizes based on token count rather than the number of proteins (Figure S2). To efficiently handle batches with varying protein lengths and maximize GPU utilization, we propose a novel methodology called Partition-Attention, which concatenates variable-length protein sequences into a single long chain without padding and utilizes attention masks to preserve batch structure (Methods). Regardless of model size, FlashAttention with Partition-Attention can handle up to 100,000 tokens without memory errors (Figure S3). This combination ensures consistent computational load across batches, achieving full GPU utilization and enhancing model inference performance 16-fold (Figure S4). The small footprint of FlashAttention and extending batch sizes using Partition-Attention could further enable training large PLMs on tasks involving protein complexes [47] and even genome-scale protein predictions [48].

We explored weight quantization techniques to optimize inference by benchmarking 8-bit quantization with LLM.int8() [49] and 4-bit quantization using QLoRA [50] algorithms using bitsandbytes package [51]. Quantization provides marginal benefits for only billion-parameter models (3B and 15B), primarily by reducing memory usage through compressing model weights to lower precision (Figure 2C, Table S2). For instance, 4-bit and 8-bit quantization reduces memory usage to 41% and 80% of the unquantized inference for 3B models and 33% and 63% for 15B models. However, the memory overhead from the quantization constants needed for quantization and dequantization limits the effectiveness of these algorithms for sub-billion-parameter models. Additionally, 8-bit quantization introduces computational overhead, nearly doubling inference time, whereas 4-bit quantization achieves a runtime comparable to bfloat16 precision (Figure S5, Table S1).

### Variant Effect Prediction

To validate the correctness of our reimplementation, we benchmarked the ESM family—ESM1v, ESM1b, and ESM2—by evaluating how effectively they capture the impact of missense mutations on protein fitness, a key application of PLMs. We used deep mutational scanning (DMS) experimental data from the ProteinGym database [52] as ground truth for 94 human proteins [52]. The ProteinGym database contains 12 proteins longer than 1024 residues, but the original ESM1b and ESM1v models use learned positional embeddings [53] with a maximum sequence length of 1024 tokens, requiring sequence truncation for longer proteins. To address this limitation in our implementation, we extended the supported sequence length to 4096 residues in the ESME-1b and ESME-1v models by retraining the positional embedding layer using longer sequences from UniRef50 [45] (Methods). Spearman’s rank correlation coefficients between predicted mask margin scores and DMS scores (Methods) demonstrate that our reimplementation produced results comparable to the original implementation on the variant effect prediction task (Figure 3, Table S3). Weight quantization leads to only a slight performance decrease with 4-bit quantization in smaller models (8M and 35M), while larger models are robust to quantization. Interestingly, model performance improves as the parameter size increases, reaching a peak at 650 million parameters for ESM2, ESM1b, and ESM1v models, with average performance of 46.6 ± 17.5%, 42.7 ± 17.2%, and 43.1 ± 16.7%, respectively. However, performance does not increase further for the larger 3 billion and 15 billion parameter models, staying at 45.2 ± 16.5% and 44.1 ± 16.6%, respectively. This suggests that the 650 million parameter model is optimal for variant effect prediction, and the reduced performance of larger ESM2 models may result from overfitting, potentially leading to an overestimation of the reference amino acids’ probabilities in the masked margin score.

**Figure 3:**
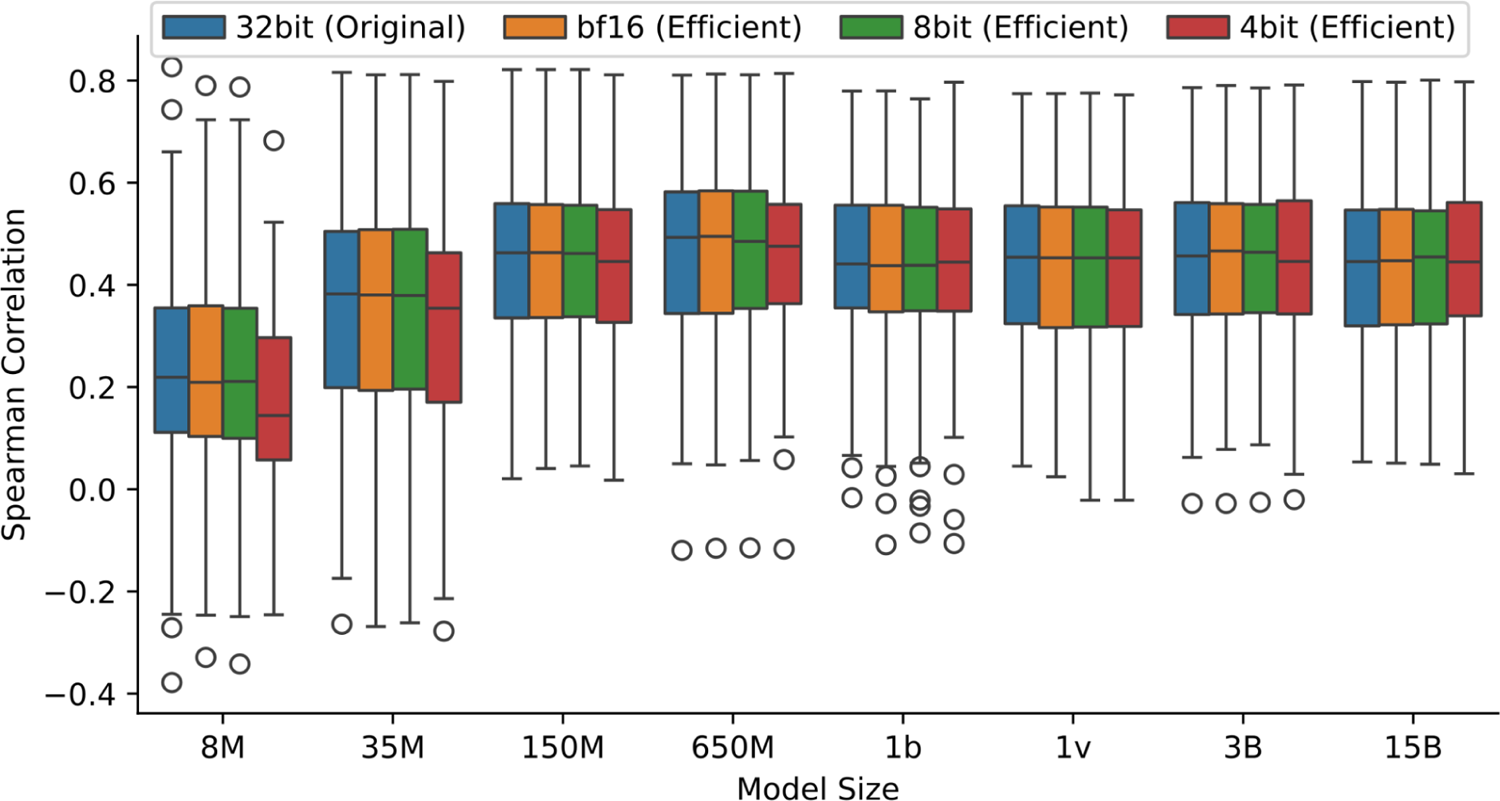
Benchmark of protein language models for predicting deep mutational screening (DMS) ground truth. The figure shows the distribution of Spearman correlation coefficients between the predicted masked margin scores by the protein language models and the DMS measurements for original implementation with full precision (blue), efficient implementation bfloat16 precision (orange), 8-bit quantization (green) and 4-bit quantization (red). The 650 million parameter models demonstrate the best performance for variant effect prediction. Quantized 8-bit and 4-bit models perform similarly to unquantized models, except for models with fewer than 150 million parameters, where 4-bit quantization results in reduced performance.

### Benchmarking the Model Training and Fine-tuning

We also benchmarked the performance of protein language models during training and fine-tuning. We measured memory savings by progressively applying various optimization techniques (Figure 4A, Table S4). Similar to inference, FlashAttention resulted in substantial memory reductions during training. However, training demands more memory than inference due to gradient computation, and Adam optimization states [54]. Therefore, we integrated training-specific techniques for backpropagation and parameter optimization to further reduce memory consumption. First, activation checkpointing reduces memory usage by recalculating intermediate states during the backward pass instead of storing them, trading runtime for memory savings, thus lowering GPU memory usage from ∼10 GB to 8 GB for the 650 million parameter model and from ∼35 GB to ∼28 GB for the 3 billion parameter ESME models.

**Figure 4:**
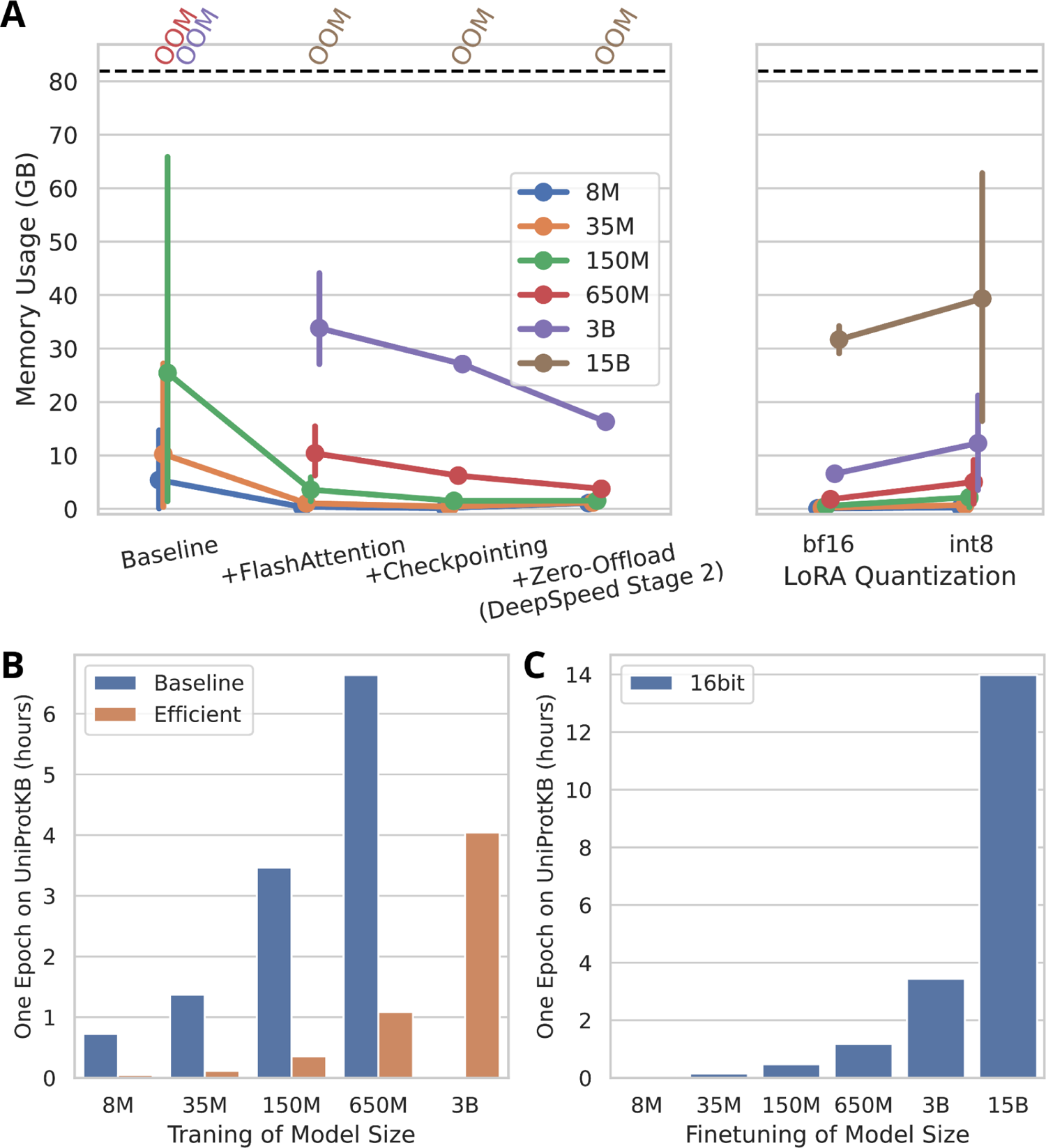
The benchmark of training performance of ESME-2. **(A)** Memory utilization of model training when the range of training and fine-tuning optimization methods are applied to the ESME-2. OOM indicates an out-of-memory error. **(B)** One epoch training time of the original ESM2 models and the efficient implementation on the UniProtKB/Swiss-Prot dataset. **(C)** The runtime of parameter efficient fine-tuning of ESME-2 models for one epoch on UniProtKB/Swiss-Prot.

Another technique, ZeroOffload/CPUAdam, offloads optimization states to CPU memory and performs parameter updates on the CPU (Methods) [55]. This strategy saves ∼39% memory for both the 650 million and 3 billion parameter models. Yet, due to memory limitations, training 15 billion parameters is still not feasible even with all the optimization algorithms with a batch size of 4 proteins.

We measured memory consumption during parameter-efficient fine-tuning by adding LoRA [56] adapters with a rank of 16 to the query, value, and output layers (Figure 4A). Fine-tuning with these adapters updates only a tiny fraction of the weights, enabling fine-tuning of the 15 billion parameter model (Methods). The largest 15 billion parameter model uses 33 GB of memory, while the remaining models use less than 10 GB. While quantizing frozen model weights during fine-tuning is possible, the current implementations of 8-bit and 4-bit quantization result in increased memory usage as the input token count grows (Figure S4). For example, quantized LoRA fine-tuning with 8-bit quantization increases GPU memory usage from ∼32 GB to ∼40 GB. Thus, unlike inference, quantization during fine-tuning is not memory efficient, highlighting the need for more effective quantization methods for PLMs. Moreover, ZeRO-Offload/CPU-Adam does not significantly reduce memory usage during parameter-efficient fine-tuning, as only the adapters are optimized; thus, offloading optimization states and gradients to CPU memory provide negligible GPU memory savings (Figure S7).

Moreover, we benchmarked the training runtime of ESME-2 after applying all the outlined optimization methods and compared it to the original implementation (Figure 4B, Table S5). Significant speedups were achieved; for instance, one epoch on 571,609 proteins from UniProtKB/Swiss-Prot takes ∼0.04, ∼1.1, and ∼3.4 hours for the 8 million, 650 million, and 3 billion parameter models, respectively, using 4 A6000 GPUs. We also estimated the training time for ESM2 models on UniRef50 (65,672,468 proteins, totaling ∼18.7 billion tokens) if trained from scratch with our implementation and setup. The original training runs for 500,000 steps with a batch size of 2 million tokens, translating to approximately 53 epochs on UniRef50. Based on this estimate, training the 150 million and 650 million parameter models would take 33 and 309 days, respectively. Lastly, we measured the runtime of fine-tuning and observed that LoRA fine-tuning is slightly faster compared to the training of all weights (Figure 4C). For instance, fine-tuning 3 billion parameter models takes ∼3.4 hours, which is 12.5% faster.

### Protein Melting Point Prediction

Annotation of protein function and properties is only available for a limited number of proteins. Thus, leveraging semi-supervised protein language models (PLMs) as foundation models to train task-specific models with limited data is critical. Semi-supervised training of PLMs on diverse sequences enables the capture of latent spaces and protein properties, allowing the model to generalize from limited training samples. To demonstrate this, we fine-tuned ESME-2 using head-only and LoRA fine-tuning to predict the melting point [57] of proteins based on their sequences (Methods). Based on the hyperparameter tuning, adapters with a rank-size of 16 were added to the query, value, and output layers for each transformer block. We achieved a Spearman’s correlation coefficient of ∼70% using head-only fine-tuning of the ESME-2 3B model for melting point prediction, surpassing the state-of-the-art [26] (Figure 5A, Table S6). LoRA fine-tuning showed improvements for smaller models, boosting performance from 58% to 62% for the 8M model and from 60% to 64% for the 35M model (Figure 5B). The test set consisted of proteins with less than 20% sequence identity to the training set, demonstrating that PLMs can generalize effectively to proteins from unrelated families or remote homologs when predicting protein properties.

**Figure 5:**
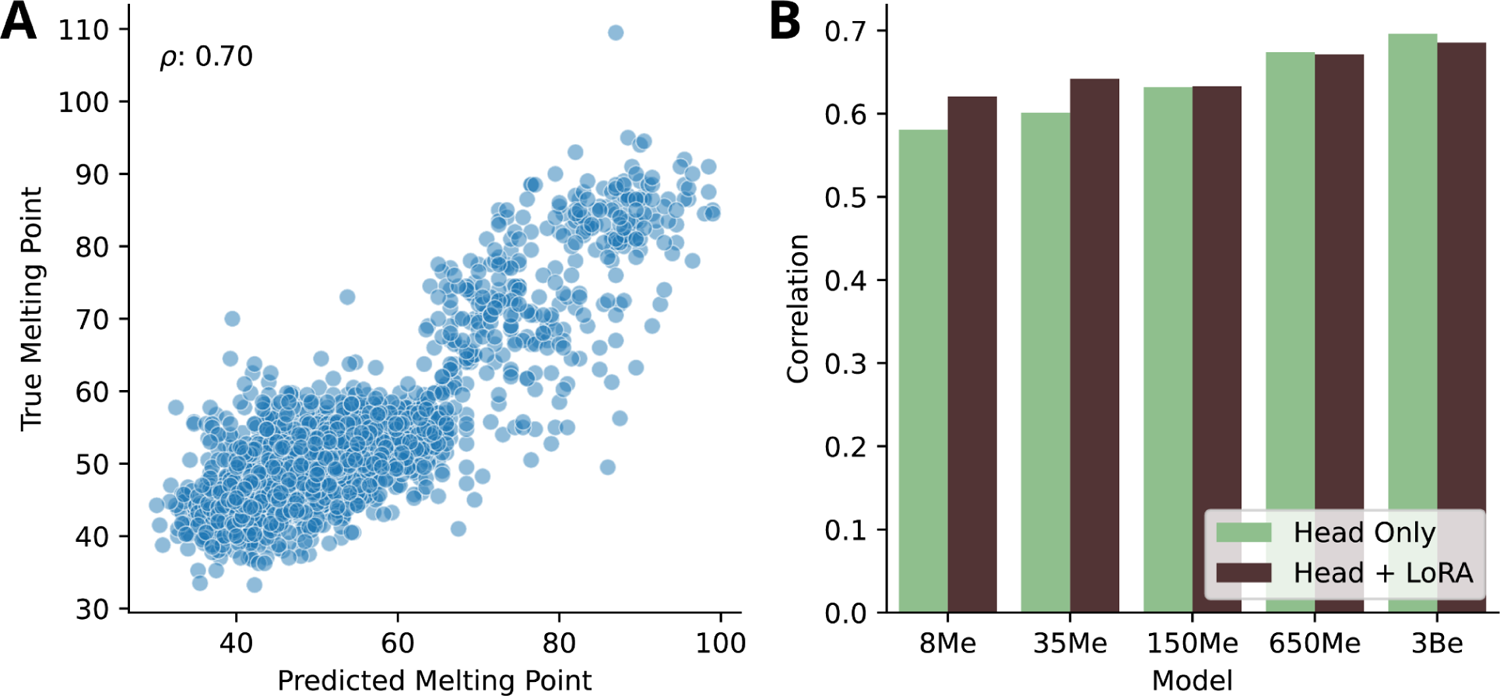
Melting Point Prediction with parameter efficient fine-tuning. **(A)** Correlation between experimentally measured protein melting points from the Meltome Atlas and predictions from the ESME-2 3B model using head-only fine-tuning. **(B)** Spearman’s correlation coefficients between observed and predicted melting points, based on head-only (green) and LoRA fine-tuning (brown) of ESME-2 models with different parameter sizes.

### Homology-based Transcription Factor Prediction

Moreover, we fine-tune the ESME-2 model to predict transcription factors (TFs) using ground truths derived from TF-related GO-terms (Methods). The train and test sets are separated by release time in the database, following the same approach as DeepTFactor. Thus, the prediction task primarily uses homology modeling to estimate protein function. The head-only fine-tuning achieves an area under the receiver operating characteristic curve (AU-ROC) of 97% on the test set, outperforming the state-of-the-art DeepTFactor model [58], which achieves 87% AU-ROC (Figure 6A, B, Table S7). The LoRA fine-tuning with a rank-size of 16, applied to the query, value, and output layers, achieves slightly higher precision, 87% under the area of the precision-recall curve (AU-PRC) for the 3 billion parameters model, while head-only fine-tuning achieves 84% AU-PRC (Figure 6C, D, Table S8). These results demonstrate that protein language models significantly enhance homology-based protein function prediction through a simple fine-tuning procedure.

**Figure 6:**
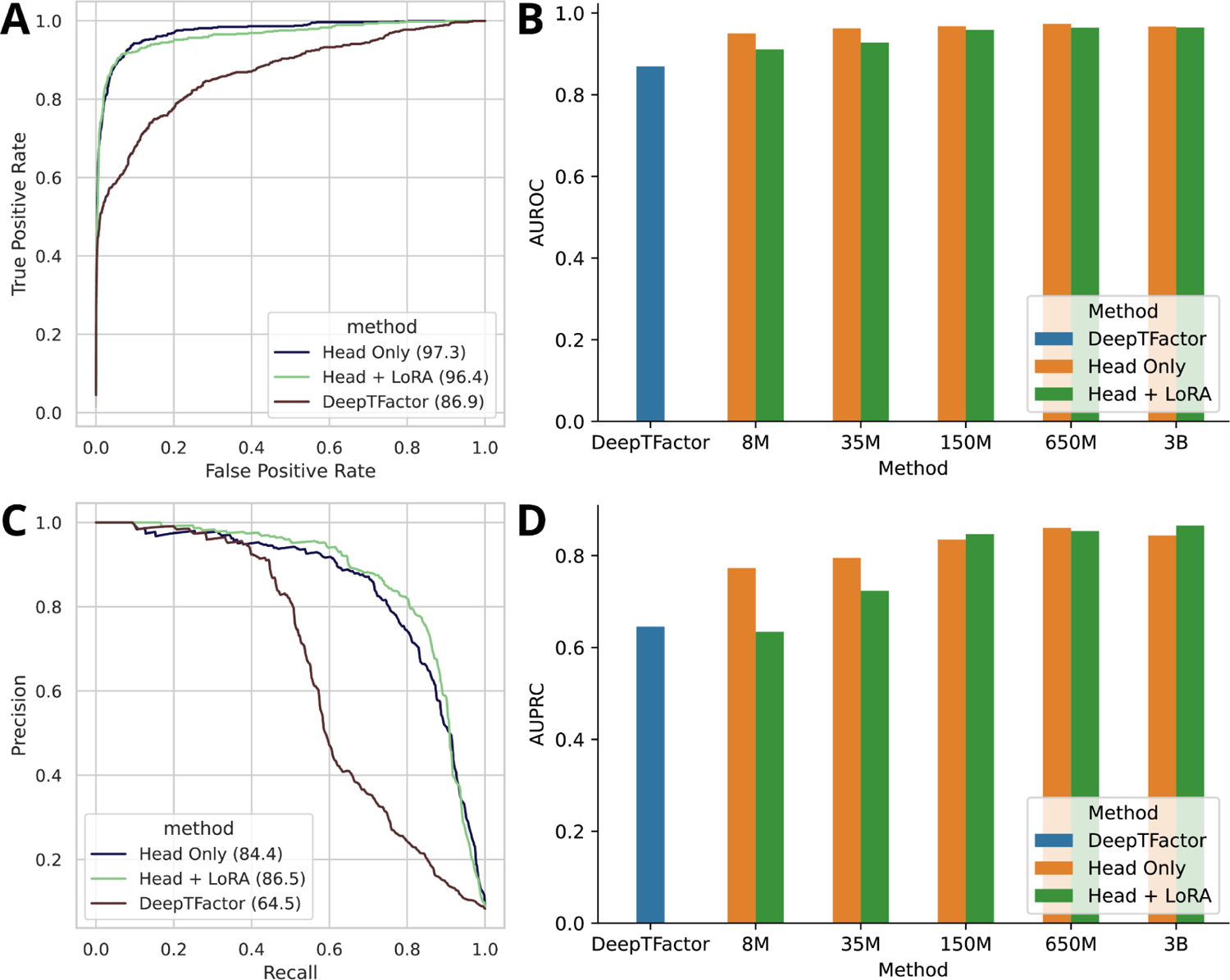
Transcription factor prediction with parameter efficient fine-tuning. **(A)** The Receiver operating characteristic curve (ROC) for transcription factor prediction of DeepTFactor, head-only, and LoRA fine-tuned ESME-2 650 million parameter model. **(B)** The area under the ROC (AUROC) for DeepTFactor and head-only and LoRA finetuned ESME-2 models. **(C)** The precision-recall curve (PRC) for transcription factor prediction of DeepTFactor, head-only, and LoRA fine-tuned ESME-2 3 billion parameters model. **(D)** The area under the ROC (AU-PRC) for DeepTFactor and head-only and LoRA finetuned ESME-2 models.

## Discussion

In this study, significant efficiency improvements are demonstrated in the training, inference, and fine-tuning of protein language models using a combination of advanced techniques such as FlashAttention, activation checkpointing, zero-offload, and low-rank adaptation. Additionally, we introduced Partition-Attention, an innovative technique that enables transformer models to effectively handle proteins with highly diverse lengths while also significantly extending the batch size. Our efficient reimplementation of the evolutionary sequence model (ESME) achieved up to a 16-fold reduction in inference time and an 8-fold reduction in memory usage. Moreover, the correctness of our reimplementation is validated by comparing it with the original model on the missense variant effect prediction task, using deep mutational scanning data as the ground truth for benchmarking. These enhancements enable the use of large-scale protein language models in more resource-constrained environments, such as academic labs.

Our findings underscore the practical utility of these models in various biological contexts. Specifically, the application of protein language models to predict protein properties and functions—such as melting points and transcription factors—demonstrates their effectiveness.

For instance, fine-tuning ESME-2 for the transcription factor prediction task achieved state-of-the-art performance while training only a small fraction of the model parameters. The LoRA fine-tuning method achieved slightly higher performance, with an AU-PRC of 87%, compared to head-only fine-tuning on transcription factor prediction. In melting point prediction, head-only fine-tuning attained the best performance, with a Spearman’s correlation coefficient of 70%, while LoRA fine-tuning was beneficial only for smaller model sizes. These results suggest that LoRA fine-tuning might achieve better performance than head-only fine-tuning on certain tasks; therefore, it should be considered another alternative fine-tuning option. Overall, the LoRA adapter weights, trained from a small amount of labeled data, can generate annotations of unannotated protein sequences via homology modeling or de novo predictions. Moreover, the adapter weights are highly modular, allowing different adapters to be easily integrated into the base models.

Our observations indicate that quantization techniques effectively save memory by compressing model weights during inference while maintaining comparable results in variant effect prediction. However, their efficacy is limited to models with billions of parameters, and the llm.int8() algorithm doubles the inference runtime, rendering it impractical. Moreover, the open-source implementation of quantization algorithms during fine-tuning can result in increased memory usage, highlighting the importance of benchmarking before adopting any optimization technique. There are quantization techniques that combine knowledge distillation with quantization [59], [60]. Although this technique might improve the model performance, further systematic benchmarking of quantization methods is another research avenue beyond the scope of this work.

Overall, the methodologies employed in this study represent a significant advancement toward making protein language models more accessible and practical for a wider range of research applications. The techniques explored are generalizable and not specific to ESM models, and we anticipate that they will be utilized in training the next generation of protein language models. Furthermore, given that the development of optimization techniques for large language models is still an active area of research, continuous benchmarking and integration of these techniques into protein language models will be essential in the future.

## Methods

In this section, we first outlined the transformer architecture of the protein language models. We then described the combination of optimization techniques applied–such as FlashAttention, Partition-Attention, model quantization, and checkpointing–to enhance the performance of ESM2 during training, fine-tuning, and inference.

### Protein Language Models

The attention layers in the model are composed of three key components: embedding layers, transformer blocks, and the language model head. The embedding layers generate embeddings for each amino acid in the input sequence. Positional information of the amino acids is incorporated through learned positional embeddings [61] in ESM-1b and ESM-1v, while ESM2 employs rotary position embeddings [62]. The transformer layers, which contain the majority of the parameters of the models, contextualize the input embeddings by capturing interactions between amino acids. These interactions are computed using self-attention, which is formalized as follows:

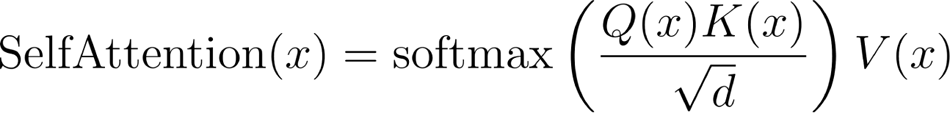

where query, key, and value vectors are used to calculate attention scores between amino acids. Each transformer block consists of an attention layer, followed by two feedforward dense layers with GeLU activation [63]. Finally, the language model head computes the logits for each amino acid based on the sequence context. The model is trained using a masked language modeling approach, where some amino acids are masked, and the task is to predict the masked amino acids using the surrounding sequence context, optimized with cross-entropy loss.

### FlashAttention

The memory footprint of standard dot-product attention grows quadratically with sequence length, creating a bottleneck when running models on long protein sequences. FlashAttention, an I/O-aware attention algorithm, addresses this issue by significantly reducing the memory footprint and improving the runtime of attention computations [44], [64]. Memory access dominates the runtime in the attention calculation. FlashAttention optimizes attention by minimizing the number of read/write operations between the slower GPU high-bandwidth memory (HBM) and the faster on-chip SRAM. Specifically, FlashAttention divides the input key, query, and value matrices into smaller blocks, loading only the necessary blocks into SRAM. All computations are then performed using fused kernels, and intermediate matrices, such as those required for the backward pass, are recalculated as needed to minimize HBM access. By fusing operations together, FlashAttention further accelerates the attention calculation. One limitation of FlashAttention is that it cannot be used for protein structure prediction because structure prediction requires calculating the full attention matrix, which is not stored during the tiled calculation.

### Partition-Attention

We propose a novel technique called Partition-Attention to efficiently handle batches of protein sequences with varying lengths. Traditionally, tokens are padded to accommodate flexible input sizes during batch preparation. However, proteins exhibit widely differing sequence lengths, which leads to batches containing a large number of padding tokens and results in wasted computational resources. To address this issue, Partition-Attention constructs a single long amino acid chain by concatenating all protein sequences in the input batch without any padding tokens and prepares an attention mask that corresponds to the original batch structure. This approach enables batching with a balanced number of tokens, ensuring consistent computational cost in each step and near-full utilization of GPUs. Furthermore, preparing batches with a consistent computational load is essential for distributed data parallelization; thus, Partition-Attention is critical for scaling computations across multiple GPUs.

### Retraining the Positional Embedding Layer

We retrained the positional embedding layers of the ESM-1v and ESM-1b models to extend their context size from 1,024 to 4,000 amino acids. To accomplish this, we constructed a training dataset by selecting proteins longer than 1,000 amino acids from the UniRef50 database, reserving 1% of the data as a validation set. During training, we froze all weights except for the positional embedding layer. We masked 15% of the input tokens and used cross-entropy as the loss function, training the model until convergence.

### Quantization

As protein language models increase in size, they can consume a significant amount of memory. Rounding model weights to lower-precision numbers can significantly reduce memory usage; for example, half-precision Brain Floating Point (bfloat16) is widely preferred over full-precision float32 due to its lower memory footprint and faster training times [46]. Model weights can even be represented with lower precision, such as 8-bit and 4-bit formats. For instance, absolute max quantization reduces a matrix to the desired precision by dividing each value by the absolute maximum value, multiplying by the quantization level, and rounding to the nearest integer. However, naive quantization methods introduce errors, thereby significantly degrading accuracy. Therefore, quantization methods specific to language models have been proposed to ensure model performance while reducing memory footprint and runtime. These quantization techniques are helpful during model inference by reducing memory usage and runtime. However, training with quantized weights may not be stable. Nevertheless, quantization can be used during fine-tuning because the quantized model weights are frozen, and only the fine-tuned weights are trained without quantization.

### 8-bit Quantization with LLM.int8

It has been empirically observed that the weights of language models are normally distributed and centered around zero. However, rare outlier features with very large magnitudes can occur. These outlier features are sufficient to shift the quantization scale, thereby introducing large errors in matrix multiplication. To address this issue, LLM.int8 [49] proposes a mixed-precision decomposition methodology that retains outlier features in the original float16 precision and performs calculations in half-precision, while computations for the remaining non-outlier weights are carried out in 8-bit using absolute max quantization. Since the outliers are extremely sparse, storing them in half-precision does not significantly affect the memory footprint while maintaining model performance.

### 4-bit quantization with QLoRA

QLoRA is an algorithm that efficiently fine-tunes large language models using 4-bit quantization. It employs blockwise quantization, where model weights are divided into blocks that are quantized independently [50], effectively containing outliers within each block. This technique stores the quantized weights using the information-theoretically optimal NormalFloat (NF) data type and further reduces memory usage by quantizing the quantization constants through a proposed double quantization method.

### Activation Checkpointing

Model training is significantly more computationally expensive than model inference because the outputs of each layer and the intermediate states needed for backpropagation must be stored during the forward pass to calculate gradients during the backward pass. The accumulation of these intermediate states leads to substantial memory usage in large deep-learning models. Activation checkpointing addresses this issue by significantly reducing GPU memory usage during model training. In this approach, intermediate results are not stored in GPU memory during the forward pass but are recomputed lazily during the backward pass. As a result, activation checkpointing saves memory at the cost of additional computational time.

### Zero-Offload and CPU-Adam

Zero-Offload is a model training strategy that employs CPU offloading to reduce GPU memory usage without significantly degrading model performance [55]. In the Stage-2 Zero-Offload strategy utilized in this work, the model weights are kept on the GPU, but gradients and optimizer states are offloaded to the CPU after the execution of each layer. Model training requires storing gradients for each parameter and optimization state, such as momentum parameters for the Adam algorithm, thereby multiplying the memory usage per weight. Storing the gradients and optimizer states on the CPU saves GPU memory. However, the offloading process can lead to a slowdown of training because updating the model weights requires transferring gradients and optimizer states between the CPU and GPU. To mitigate the inefficiencies introduced by CPU-GPU communication, Zero-Offload utilizes CPU-Adam [55], an algorithm optimized for computation on the CPU. CPU-Adam performs the weight updates on the CPU based on the gradients and optimizer states stored in CPU memory and then transfers the updated model parameters back to the GPU. When this approach is combined with gradient accumulation, where gradients are accumulated over a certain number of steps before updating the model parameters, it minimizes CPU-GPU memory transfer overhead, resulting in minimal efficiency loss during model training.

### LoRA

Fine-tuning large language models (LLMs) is computationally expensive due to their large parameter sizes. Additionally, information learned during foundational model training might be lost during updates during fine-tuning, a phenomenon known as catastrophic forgetting.

Low-Rank Adaptation (LoRA) [56] addresses these issues by training only a small fraction of the model’s weights, called adapters, during fine-tuning. These low-rank adapters are added to the key, query, and value matrices and are defined as:

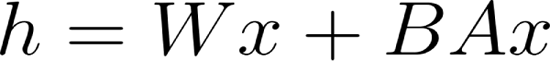

where *W ∈ R^n × m^* and *B ∈ R^n × r^, A ∈ R^r × m^* the matrix *r* rank of the matrixes *A* and *B* are much smaller (*n ≫ r,m ≫ r*) than the size of the key, query, and value matrixes. As a result, model weights are frozen, and only the extra adapter weight is trained with LoRA, leading to efficient model training and avoiding catastrophic forgetting. The adapters can be attached and detached from the model, enabling modularity.

### Variant Effect Prediction

The protein language models can prioritize potentially pathogenic missense variants by comparing the posterior likelihood of the alternative and to the reference amino acids:

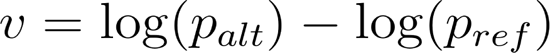

where probabilities for the reference (*p_ref_*) and alternative (*p_alt_*) alleles are predicted by the model. The deep mutational scanning (DMS) experiments provide unbiased experimental ground truth for the benchmark of variant effect prediction on protein fitness. DMS experiments for 92 human proteins were downloaded from the ProteinGym database [52] and used to validate the correctness of our reimplementation. ESM-1v and ESM-1b models only support sequences up to 1024 amino acids; thus, sequences are truncated to match this length by centering the variants. Similarly, we used this truncation strategy for the original 15 billion parameter ESM2 model to avoid memory error. For ESME models, variant effect predictions were performed with entire protein sequences.

### Melting point prediction

Protein language models can be fine-tuned to predict various properties of proteins. To demonstrate this capability, we fine-tuned models to predict the melting points of proteins. We used the Meltome Atlas [57] as the ground truth for melting points, which contains approximately 48,000 proteins from 13 species. The training and test sets were obtained from the Fitness Landscape Inference for Proteins (FLIP) project [26], which splits the data so that sequences in the training and test sets share no more than 20% sequence identity. We employed two fine-tuning approaches: head-only fine-tuning and LoRA fine-tuning. In head-only fine-tuning, we trained classification heads consisting of two feedforward layers with ReLU activation functions. In LoRA fine-tuning, additional adapters were added to the query, key, and value matrices of each transformer layer. For optimization, we used the Adam algorithm with a learning rate of 10^-4^and momentum parameters of 0.9 and 0.999. The learning rate was reduced by 10% every 10 epochs during 100 epochs of training. Mean squared error is used as a loss function.

### Transcription Factor Prediction

Another downstream application of protein language models is the prediction of protein functions. Specifically, in this work, we fine-tuned protein language models to predict transcription factors. We annotated proteins in UniProtKB/SwissProt as transcription factors if they are associated with relevant Gene Ontology (GO) terms (Table S9), using an approach adapted from DeepTFactor [58], a state-of-the-art model for transcription factor prediction. We partitioned the data into training and test sets based on the publication date of the proteins in the UniProtKB/SwissProt database: proteins published up to 2021 were used for training, while those published after this date were used for testing. We employed a fine-tuning approach similar to that used for melting point prediction, except that we used cross-entropy as the loss function and up sampled the positive class of transcription factor proteins to balance it with non-transcription factor proteins in each epoch.

## Supplementary Figures

**Figure S1:**
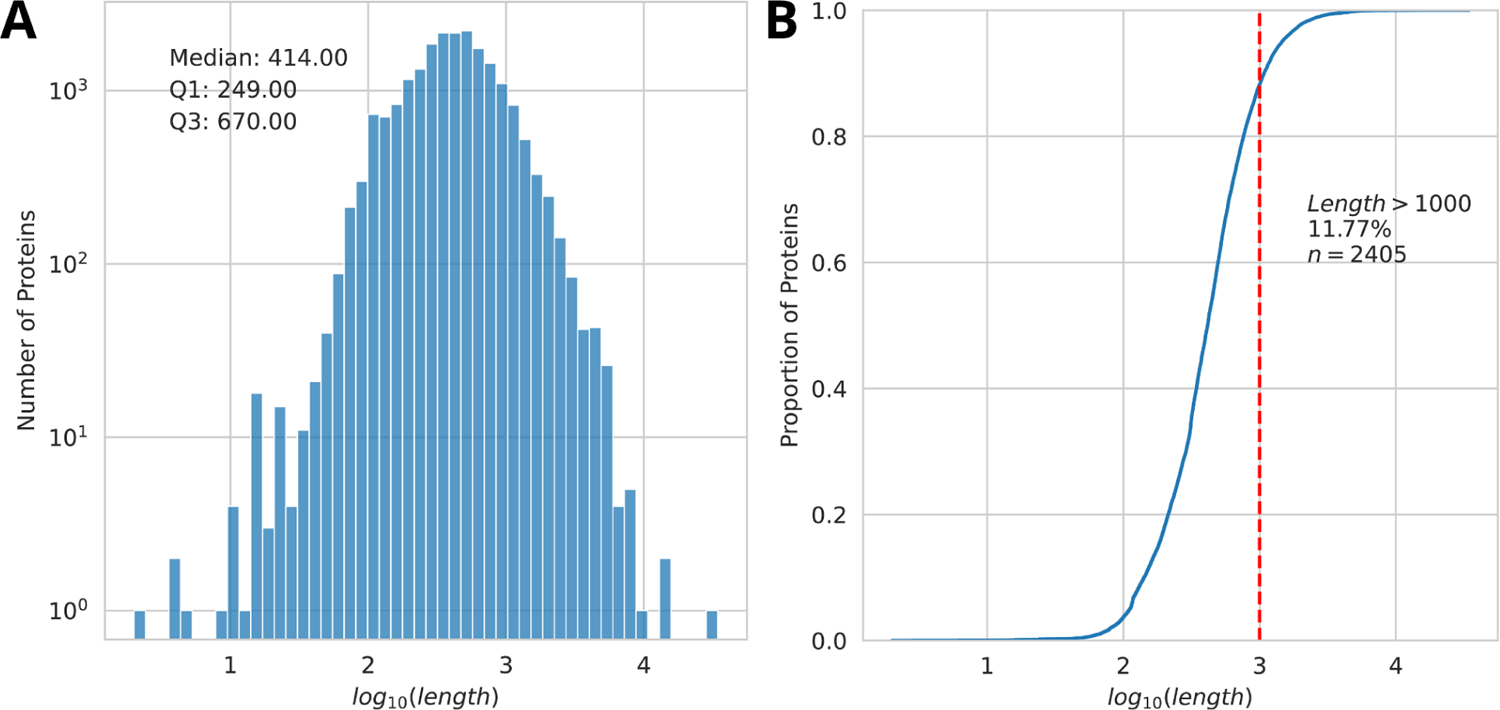
Protein Length Distribution in the Human Proteome. **(A)** The length distribution of human proteins in UniProtKB/Swiss-Prot (n=20,435). The median protein length is 414 amino acid residues. Proteins in the lower quartile (Q1) are shorter than 249 residues, while those in the upper quartile (Q3) exceed 670 residues. The longest human protein, Titin (Q8WZ42), consists of 34,350 residues. **(B)** The cumulative density distribution of protein lengths reveals that approximately 12% of human proteins (n=2,045) are longer than 1,000 residues.

**Figure S2:**
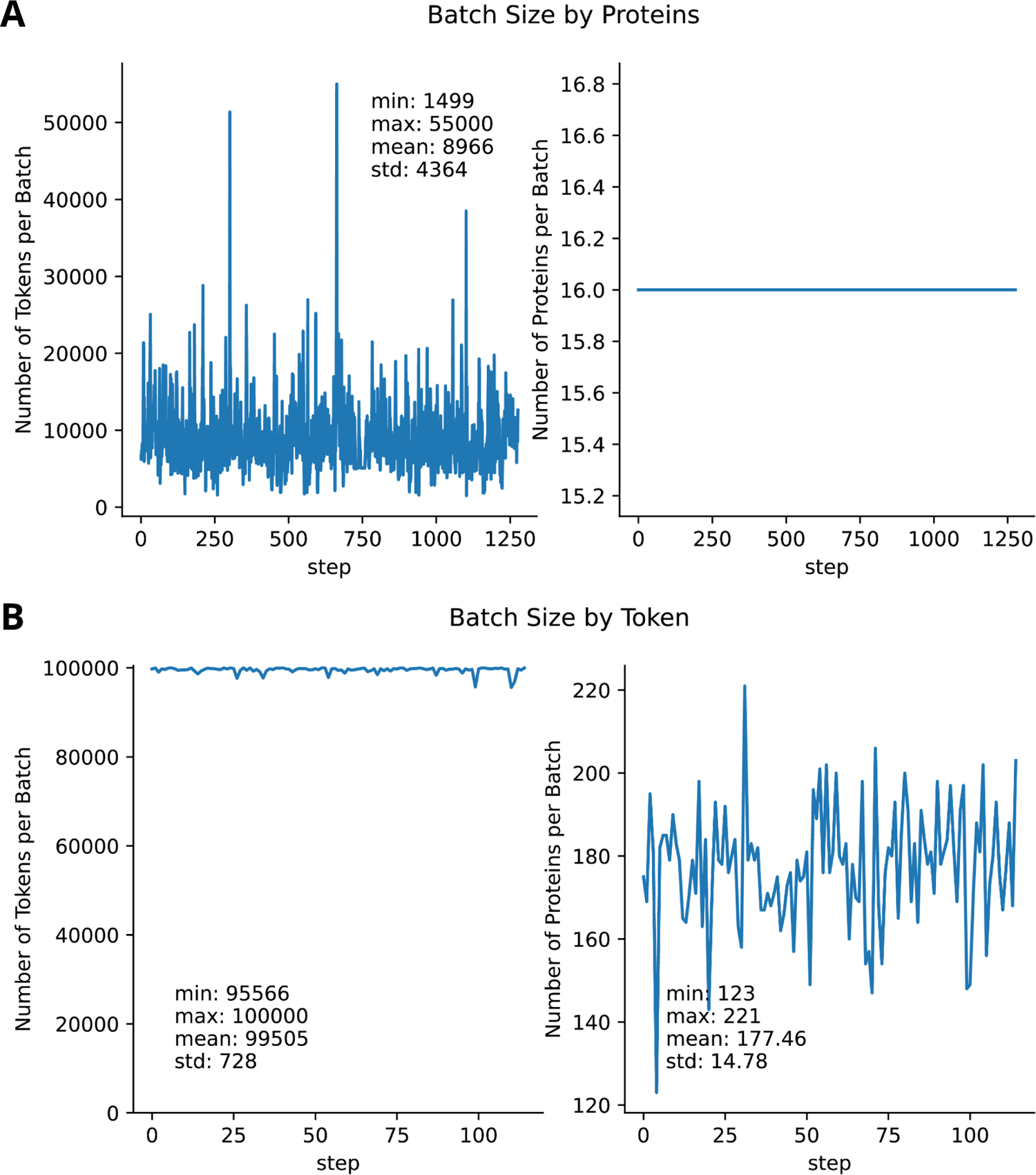
Number of Tokens in a Batch by Batching Technique. (A) The number of tokens and proteins per batch when batching is based on the number of proteins (batch size of 16 proteins). While the number of proteins per batch is constant, batch sizes vary significantly due to the variable lengths of proteins. Random batching with a fixed protein count from UniProtKB/Swiss-Prot results in an average token size of 8,966 ± 4,364. The largest batch contains 55,000 tokens, while the smallest contains 1,499 tokens. One forward pass on the dataset requires 1,266 steps. (B) Batching by the number of tokens (with a maximum of 100,000 tokens per batch) leads to more consistent batch sizes of 99,505 ± 728 tokens. However, the number of proteins per batch fluctuates due to the variation in protein lengths. One forward pass on the dataset requires only 121 steps.

**Figure S3:**
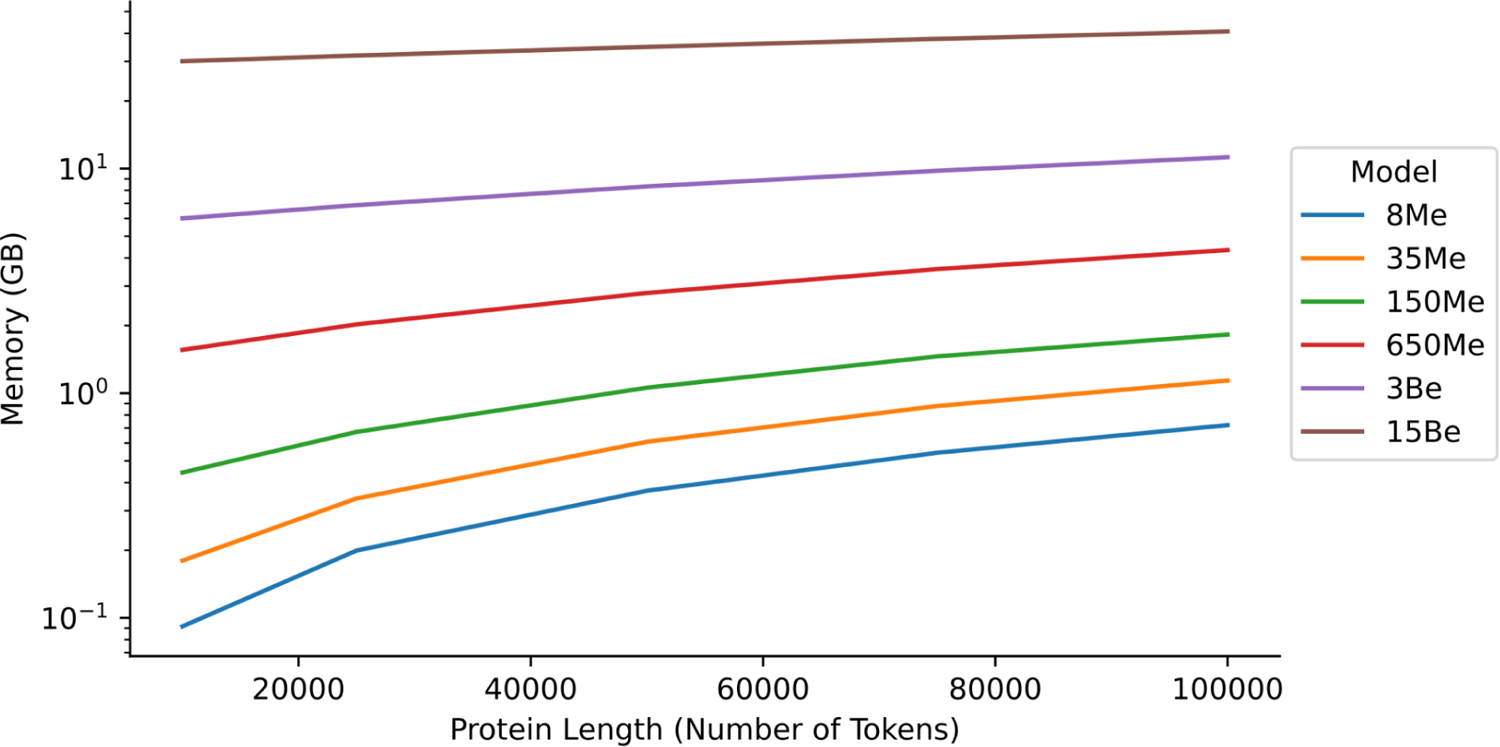
GPU Memory Usage of ESME2 Models. The sequence context can be extended to 100,000 tokens using the FlashAttention implementation in protein language models, enabling large batch sizes in conjunction with Partition-Attention.

**Figure S4:**
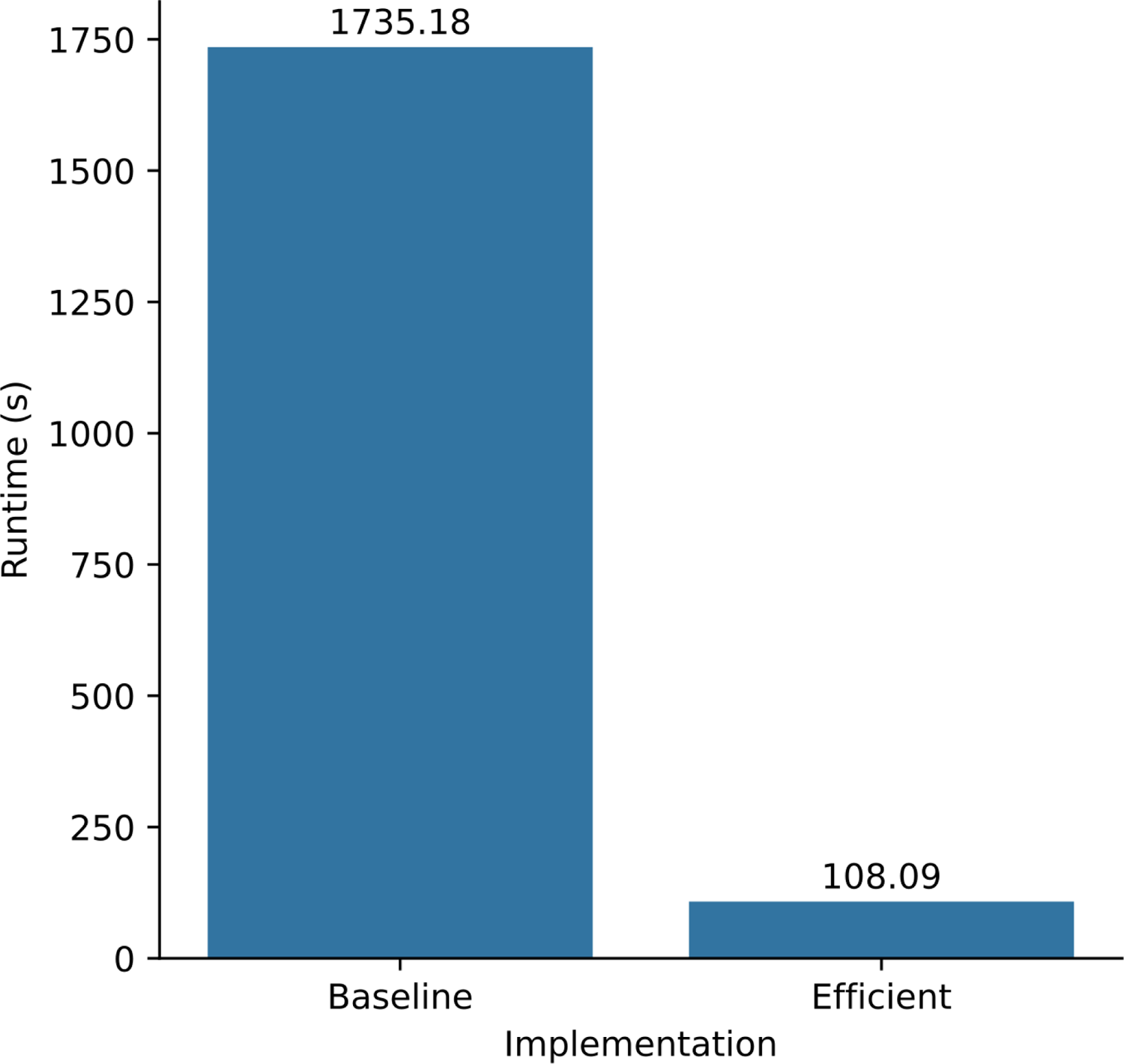
Inference Time on ESM2 Implementations. A forward pass of the ESM2 model with 650M parameters on UniProtKB/Swiss-Prot takes 1,735 seconds using the original implementation. In contrast, an optimized implementation that leverages FlashAttention and Partition-Attention reduces the inference time to just 108 seconds.

**Figure S5:**
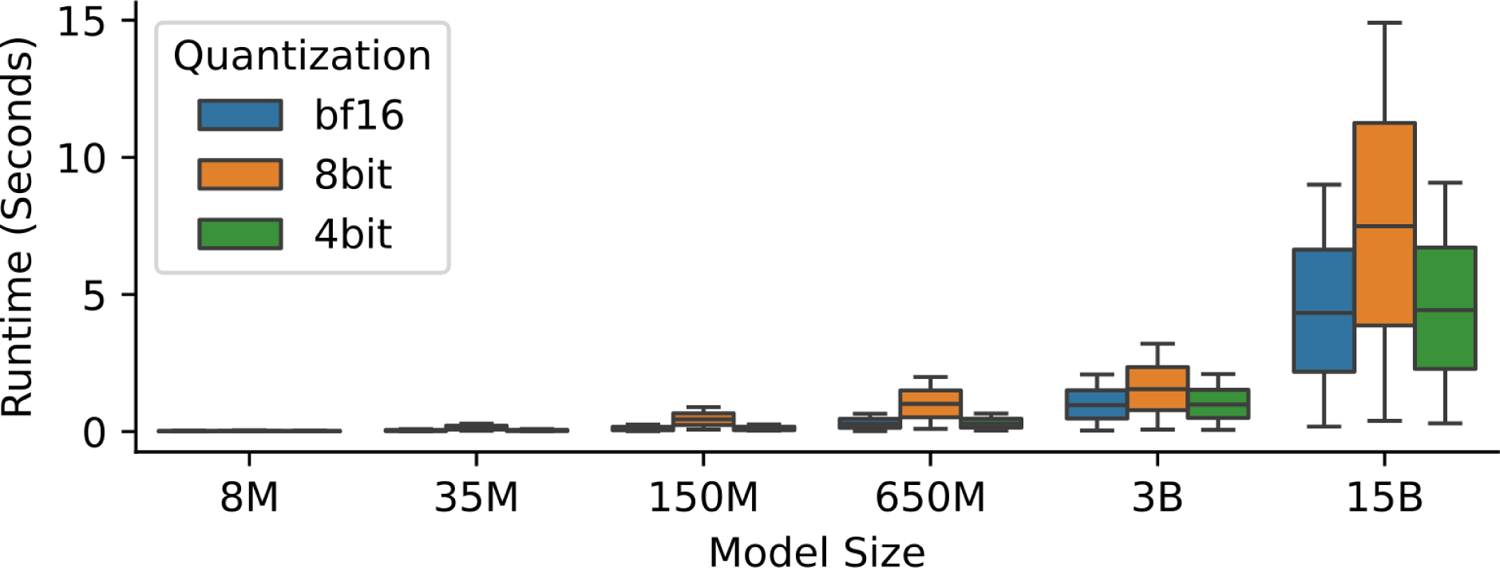
Runtime of Quantized Protein Language Models. The runtime of models using bfloat16 (brain floating point) and 4-bit quantization is comparable across different model sizes, while 8-bit quantization is slower than bfloat16.

**Figure S6:**
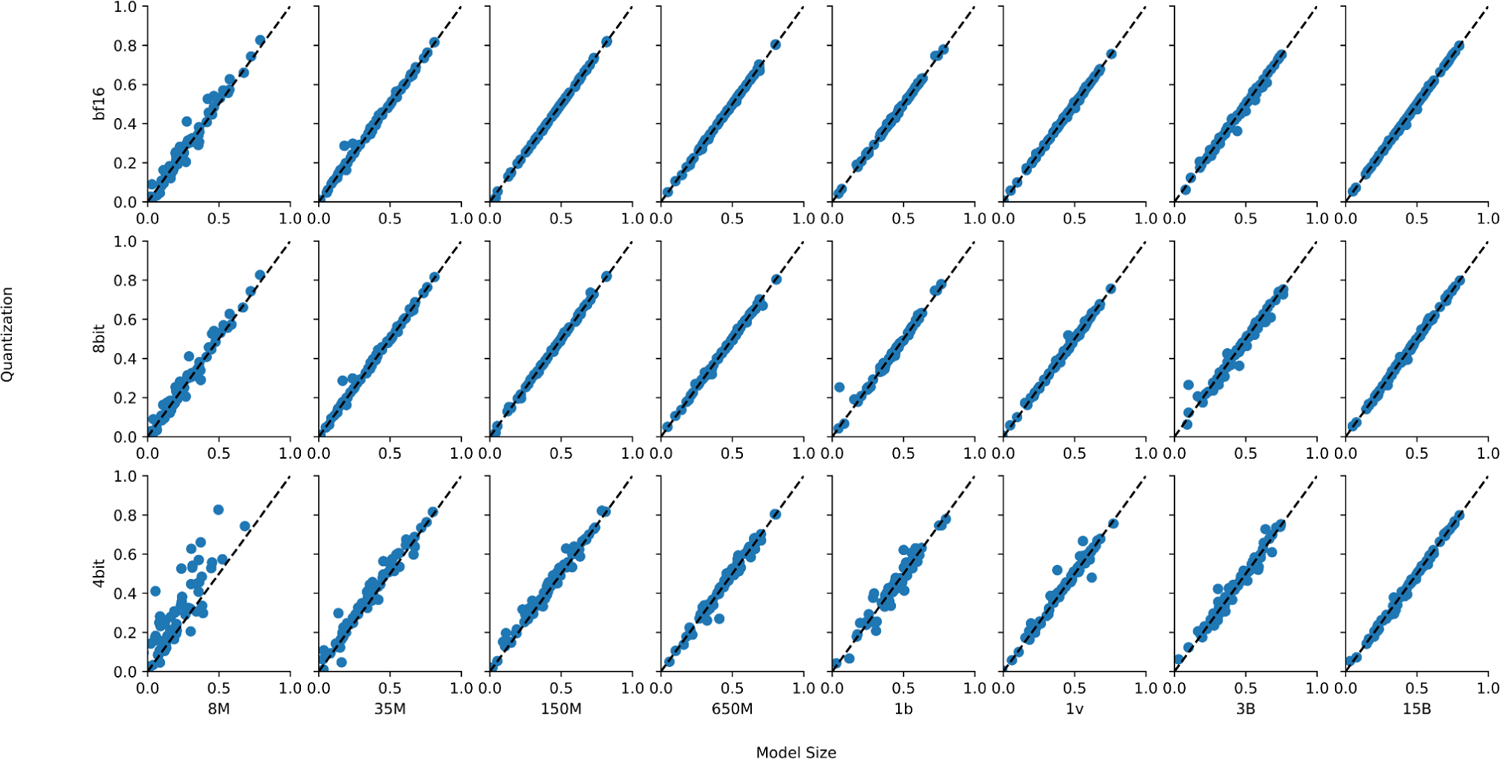
Performance of Quantized Models on Deep Mutational Screening Experiments. Lower-precision and quantized models achieve performance comparable to the full-precision (float32) model across various model sizes for variant effect prediction in deep mutational screening experiments, except for the 8 million and 35 million parameter models, where performance declines with 4-bit quantization.

**Figure S7:**
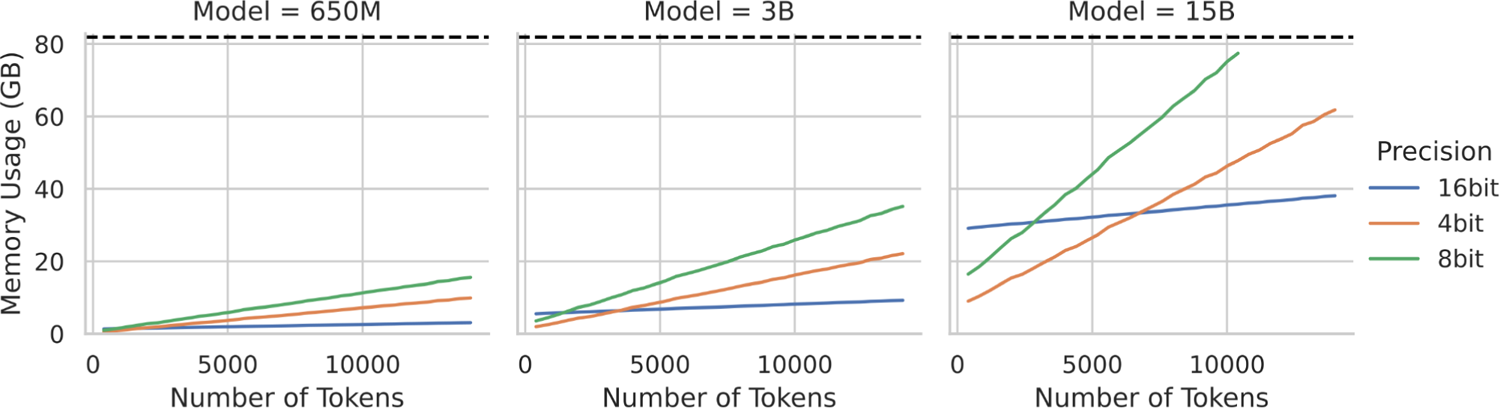
Memory usage of quantized LoRA fine-tuning. The quantized LoRA fine-tuning uses significantly more memory with increasing token sizes. The quantization of the model weights saves memory, but this saving is offset by the quantization overhead of the models.

**Figure S8:**
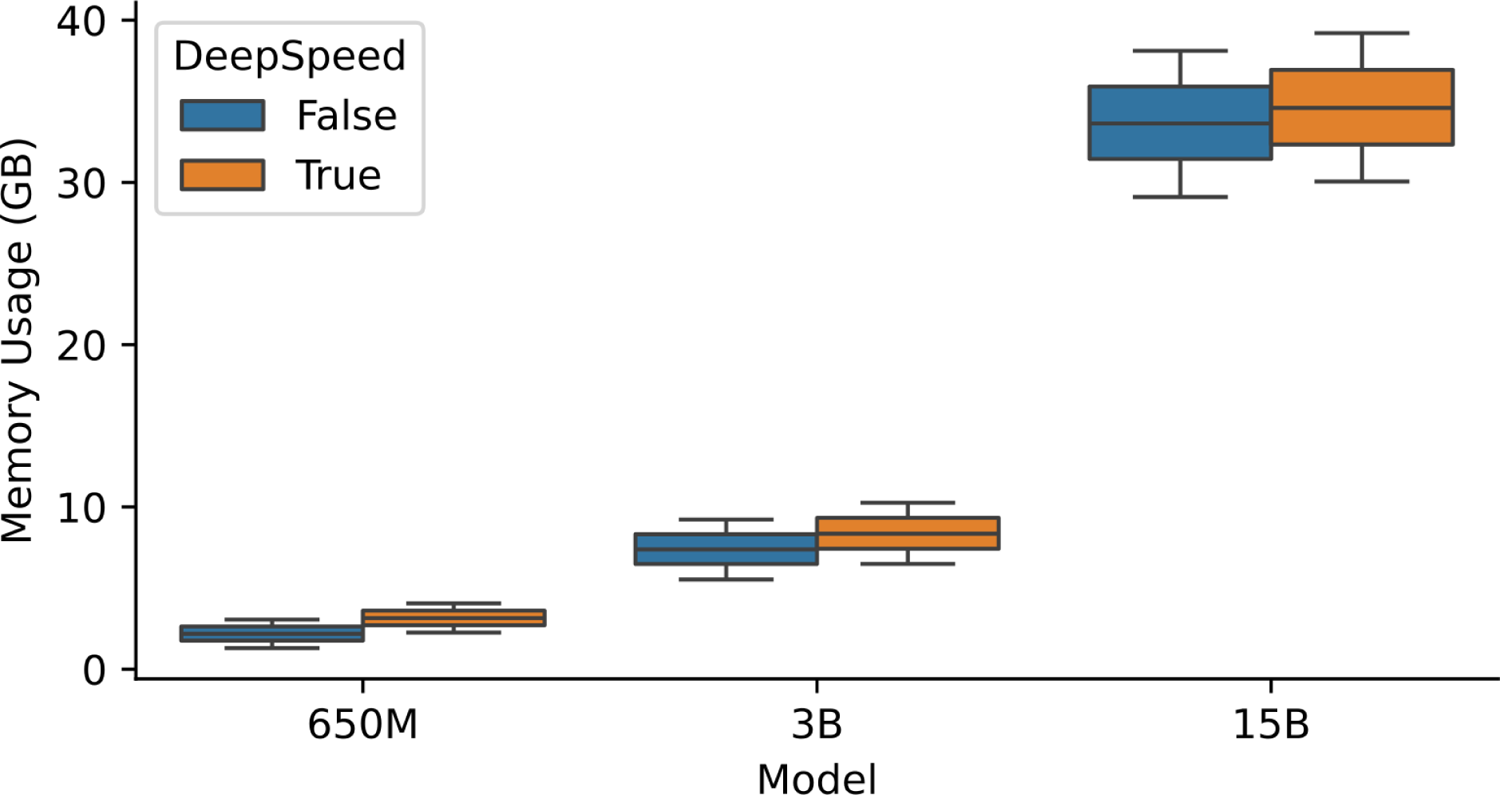
LoRA fine-tuning with/without DeepSpeed LoRA fine-tuning with DeepSpeed does not reduce memory usage, as only a small fraction of the weights have an optimization state as a result of parameter efficiency.

**Figure S9:**
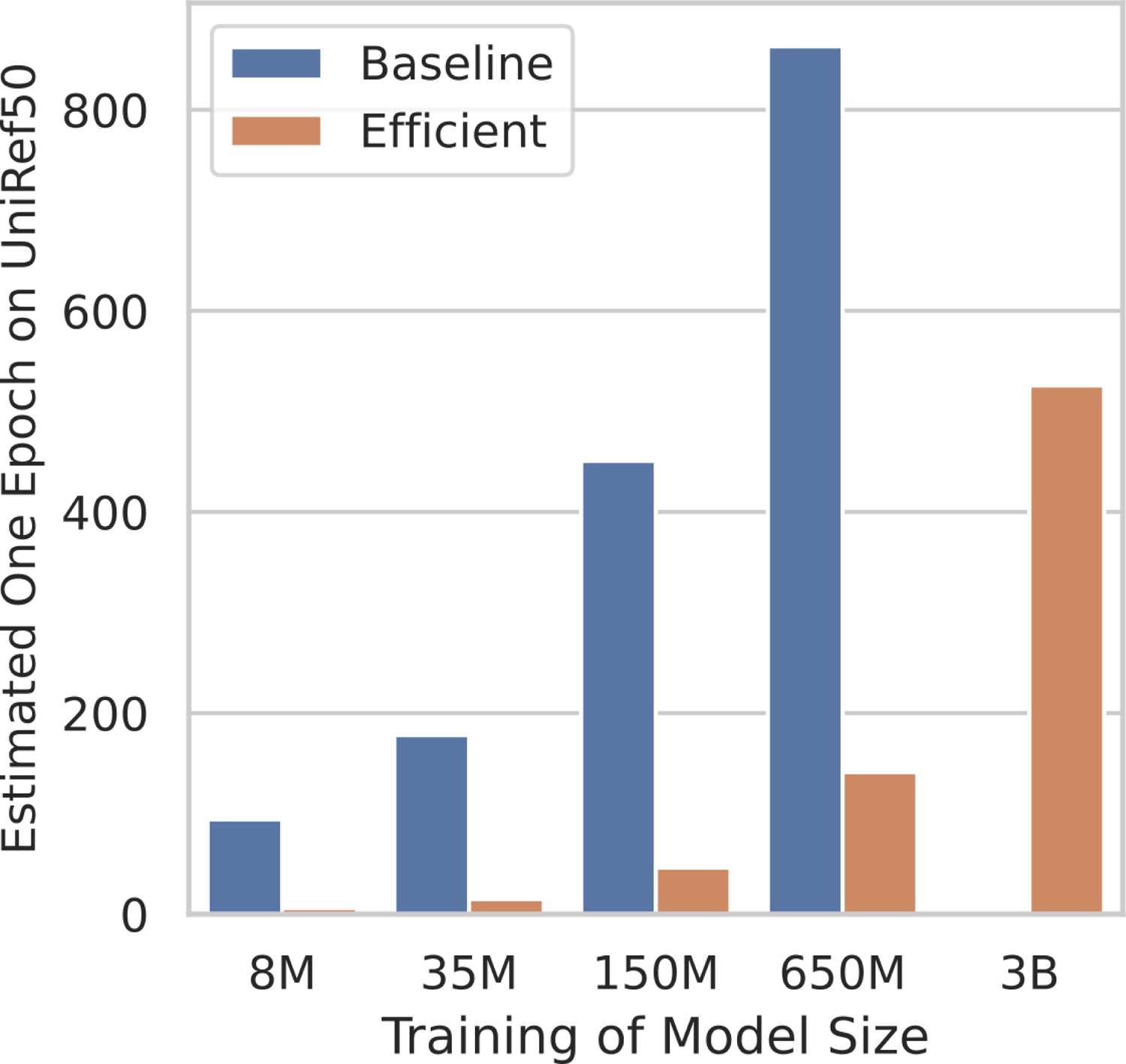
Estimated training time of the ESM2 models. Efficient implementation of the ESM2 reduces the training time of the ESM2 models 5-11 times across the model sizes. For example, one epoch on UniRef50 training of ESME-2 is estimated as ∼180 hours, while the original implementation would take ∼900 hours based on the estimated training times on 4 A6000 GPUs.

